# IKKβ as a putative non-covalent and quinone-mediated covalent target of 4-methylcatechol in RANKL/NF-κB signaling: a combined computational and experimental analysis

**DOI:** 10.64898/2026.07.29.741661

**Authors:** Chengxu Xie, Lifang Zhang, Xinyi Bao, Xiaohan Li, Yili Ding, Mojtaba Tabandeh, Farwa Basit, Heriberto Velez, Santosh Kumar, Vishwa Deepak

## Abstract

Excessive osteoclast activity contributes to pathological bone loss in osteoporosis, rheumatoid arthritis, and osteolytic malignancies. The effects of small catechol derivatives on receptor activator of nuclear factor-κB ligand (RANKL)-induced osteoclastogenesis remain poorly understood. This study investigated the effects of 4-methylcatechol (4-MC) on RANKL-induced NF-κB activation and osteoclast differentiation. 4-MC reduced RANKL-induced NF-κB luciferase activity in HEK-293T/RANK cells. 4-MC also suppressed RANKL-induced TRAP activity in RAW264.7 cells in a concentration-dependent manner and reduced the number of TRAP-positive multinucleated osteoclasts, without affecting cell viability. Molecular docking predicted non-covalent binding of 4-MC within the ATP-binding hinge region of IKKβ (PDB: 4KIK), forming a close polar contact with Glu97, predicted hydrogen bonds with Cys99, and a hydrophobic contact with Ile165, within the pocket occupied by the co-crystallized inhibitor K252a. Covalent docking predicted that the oxidized quinone form of 4-MC engages Cys179 in the IKKβ activation loop. Quantum chemical calculations confirmed a markedly higher electrophilicity index for the oxidized quinone than for the parent catechol, supporting this mechanism. In silico ADMET profiling indicated favorable drug-likeness and safety. These findings identify IKKβ as a plausible molecular target of 4-MC through both non-covalent and covalent mechanisms.

**Graphical Abstract:** 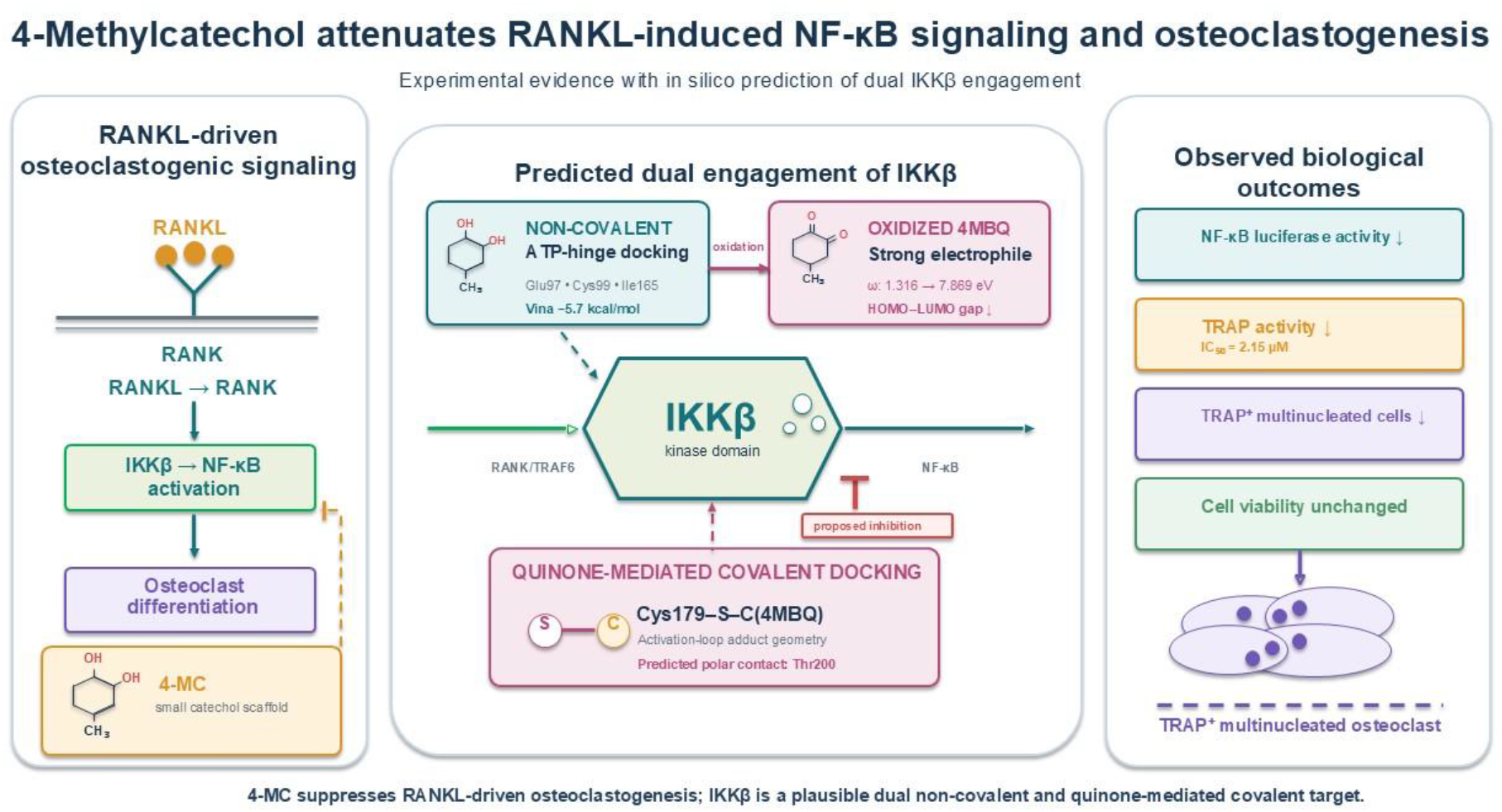

## 1. Introduction

Bone remodeling depends on coordinated osteoblast-mediated bone formation and osteoclast-mediated bone resorption. Persistent elevation of osteoclast activity shifts this balance toward bone loss and contributes to osteoporosis, inflammatory bone erosion in rheumatoid arthritis, and osteolytic malignancies (1). Osteoporosis and fragility fractures impose a substantial clinical and economic burden worldwide (2). Although bisphosphonates and RANKL-targeted biologics are effective antiresorptive treatments, their limitations support the continued search for small molecules with distinct mechanisms of action (3).

Osteoclasts are multinucleated cells derived from monocyte and macrophage lineage precursors. Their differentiation is primarily regulated by macrophage colony-stimulating factor and receptor activator of nuclear factor-κB ligand (RANKL). RANKL binding to RANK recruits tumor necrosis factor receptor-associated factor 6 and activates the IκB kinase and NF-κB pathways (4, 5). IKK activation promotes IκBα phosphorylation and degradation, allowing NF-κB subunits to translocate into the nucleus. NF- κB and c-Fos subsequently promote the induction of nuclear factor of activated T cells 1 (NFATc1), the master transcriptional regulator of osteoclast differentiation (5, 6). NFATc1 drives the expression of osteoclast-associated genes, including tartrate-resistant acid phosphatase and cathepsin K (6). The central role of NF-κB in this process makes the pathway an attractive target for suppressing excessive osteoclast differentiation.

Catechols contain two adjacent hydroxyl groups on a benzene ring. This ortho-diol motif supports hydrogen bonding and redox activity, allowing catechol-containing compounds to influence ROS-sensitive signaling pathways. Caffeic acid derivatives, including MPMCA and caffeic acid phenethyl ester, have been reported to suppress osteoclastogenesis, with CAPE inhibiting NF-κB, c-Fos, and NFATc1 signaling (7, 8). However, the effects of smaller catechol molecules without extended aromatic side chains remain poorly characterized.

4-Methylcatechol (4-MC), also known as 4-methylbenzene-1,2-diol, is a minimally substituted catechol containing a single methyl group. It retains the ortho-diol motif while introducing a small hydrophobic substituent (9). Notably, 4-MC is not merely a synthetic reference compound: it has been isolated directly from plant material (Picea abies) and also arises as a major microbial/metabolic breakdown product of rutin, a flavonoid glycoside widely distributed in dietary fruits and vegetables, placing it within a broader class of naturally occurring alkyl catechols. Previous studies identified 4-MC as a stimulator of nerve growth factor synthesis and showed that it promoted peripheral nerve regeneration in an experimental nerve-injury model (10). More recently, 4-MC was shown to suppress RANKL-induced osteoclastogenesis indirectly, via Keap1/Nrf2-dependent upregulation of heme oxygenase-1 (HO-1) and consequent reduction of intracellular reactive oxygen species (11). However, whether 4-MC additionally engages the NF-κB pathway directly, upstream or independently of this antioxidant mechanism, has not been established.

In the present study, we investigated whether 4-methylcatechol suppresses RANKL-induced NF-κB activation and osteoclast differentiation. NF-κB reporter activity, TRAP activity, and RAW264.7 cell viability were examined. Molecular docking against the active chain B of human IKKβ from PDB structure 4KIK (12) was also performed to explore IKKβ as a possible molecular target, together with covalent docking, quantum chemical reactivity analysis, and in silico ADMET profiling to characterize the compound’s dual non-covalent and covalent engagement potential.

## 2. Results and discussion

### 2.1. 4-MC attenuates RANKL-induced NF-κB activation

RANKL stimulation markedly increased NF-κB-dependent luciferase activity in HEK-293T/RANK reporter cells relative to the vehicle control (Fig. 1). Pretreatment with 4-MC (1 µM) significantly attenuated this RANKL-induced response compared with RANKL treatment alone (***P < 0.001). At the tested concentration, reporter activity remained above the vehicle-control level, indicating that 4-MC selectively dampened the RANKL-induced increase in NF-κB activity without suppressing signaling below its basal level. This reduction was not attributable to cytotoxicity, as 4-MC did not affect the viability of HEK-293T/RANK reporter cells at the tested concentration (Fig. S1).

**Fig. 1.**
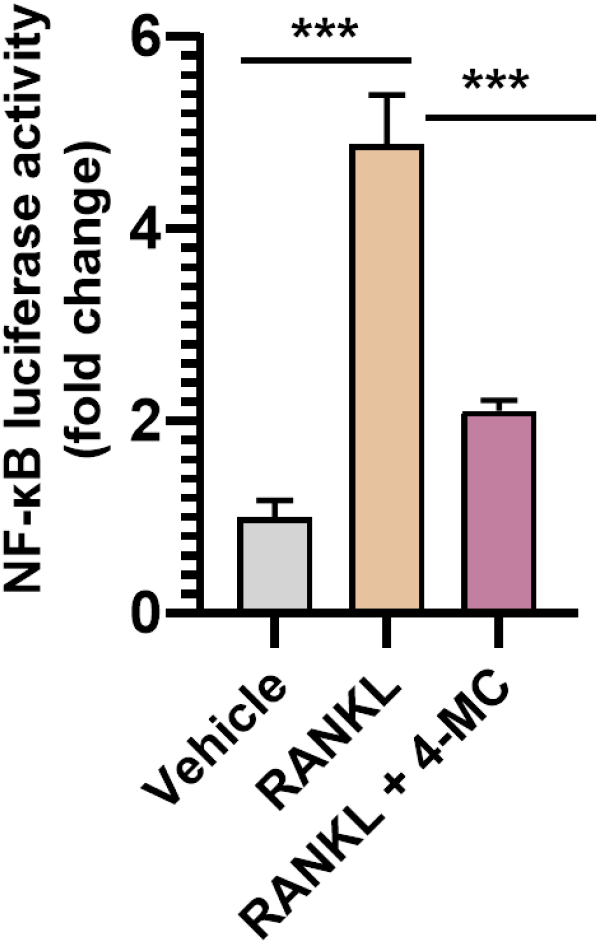
4-Methylcatechol attenuates RANKL-induced NF-κB luciferase reporter activity. HEK-293T/RANK NF-κB luciferase reporter cells were pretreated with 4-methylcatechol (4-MC; 1 µM) for 1 h and subsequently stimulated with RANKL (50 ng/mL) for 6 h. NF-κB-dependent luciferase activity was normalized to that of the vehicle-treated control and expressed as fold change. Data are presented as the mean ± SD from three independent experiments. Statistical significance was determined using one-way ANOVA followed by Tukey’s multiple-comparisons test. ***P < 0.001 for the indicated comparisons.

### 2.2. 4-MC suppresses RANKL-induced TRAP activity and osteoclast differentiation

We next examined whether attenuation of NF-κB activation by 4-MC was accompanied by suppression of the osteoclastogenic response. RAW264.7 cells were stimulated with RANKL (50 ng/mL) in the presence of increasing concentrations of 4-MC, and TRAP activity was measured colorimetrically. 4-MC inhibited RANKL-induced TRAP activity in a concentration-dependent manner. The half-maximal inhibitory concentration was determined by four-parameter logistic regression (IC50 = 2.15 µM; 95% CI, 1.821 to 2.772 µM); Fig. 2A).

**Fig. 2.**
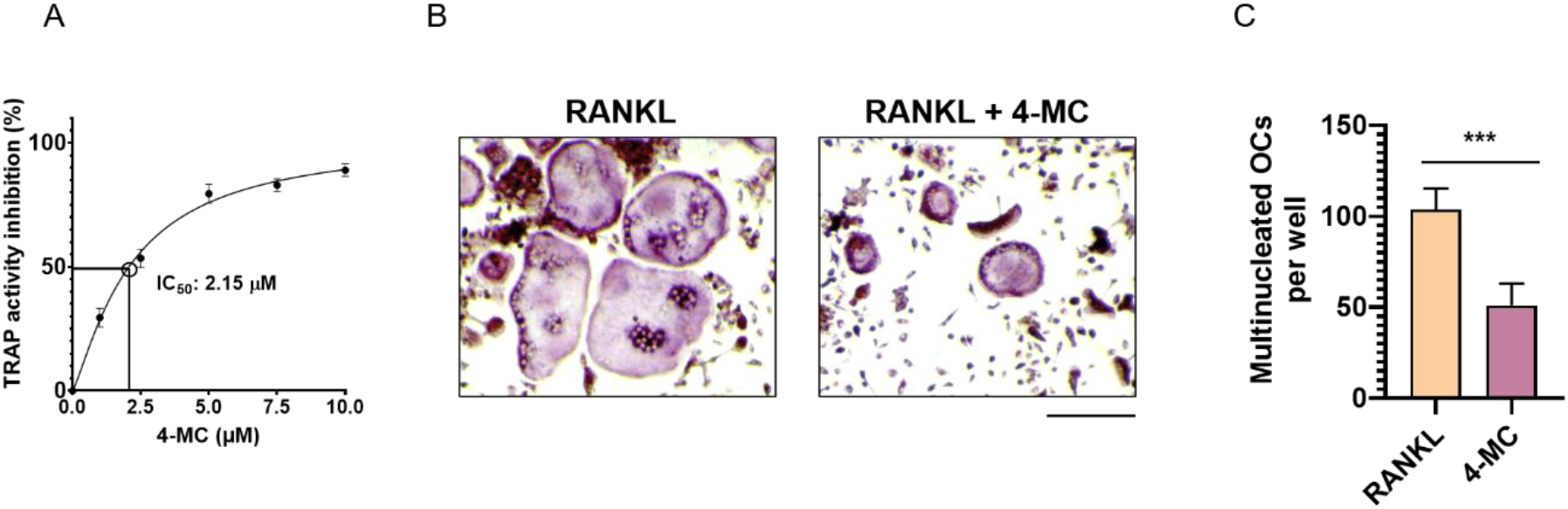
4-Methylcatechol suppresses RANKL-induced TRAP activity and osteoclast formation. (A) RAW264.7 cells were stimulated with RANKL (50 ng/mL) in the presence of increasing concentrations of 4-methylcatechol (4-MC) for 3 days. TRAP activity inhibition was calculated relative to the RANKL-treated group. The half-maximal inhibitory concentration was determined by four-parameter logistic regression (IC50 = 2.15 µM; 95% CI, 1.821–2.772 µM). Data are presented as the mean ± 95% CI from three independent experiments. (B) Representative images of TRAP-stained RAW264.7 cultures treated with RANKL alone or RANKL plus 4-MC (2 µM) for 3 days. TRAP-positive cells appear dark red or purple. Scale bar = 100 µm. (C) Quantification of TRAP-positive multinucleated osteoclasts containing three or more nuclei per well. Data in (B) and (C) are presented as the mean ± SD from three independent experiments. Statistical significance in panel C was determined using an unpaired two-tailed Student’s t-test. ***P < 0.001.

Consistent with the TRAP activity results, TRAP staining showed fewer TRAP-positive multinucleated osteoclasts in cultures treated with RANKL and 4-MC (2 µM) than in cultures treated with RANKL alone (Fig. 2B). Quantification confirmed that 4-MC significantly reduced the number of TRAP-positive multinucleated osteoclasts containing three or more nuclei (Fig. 2C). These findings demonstrate that 4-MC suppresses RANKL-induced osteoclast differentiation and formation.

NF-κB signaling is essential for osteoclast differentiation and contributes to the induction of c-Fos and NFATc1 following RANKL stimulation (5, 6). Therefore, attenuation of NF-κB transcriptional activity provides a plausible signaling basis for the observed reduction in TRAP activity and osteoclast formation. However, the reporter assay does not identify the specific signaling protein affected by 4-MC.

Importantly, 4-MC did not significantly reduce RAW264.7 cell viability at the tested concentrations (Fig. S2), indicating that its inhibitory effect on osteoclast differentiation was unlikely to result from nonspecific cytotoxicity. Caffeic acid derivatives, including MPMCA and caffeic acid phenethyl ester (CAPE), have previously been reported to suppress osteoclastogenesis, with CAPE inhibiting NF-κB, c-Fos, and NFATc1 signaling (7, 8). The present findings extend this structure-activity relationship by showing that a minimally substituted catechol scaffold lacking the extended aromatic side chains of CAPE and MPMCA can also suppress RANKL-induced NF-κB activation and osteoclast differentiation.

### 2.3. Molecular docking predicts non-covalent binding of 4-MC within the IKKβ ATP-binding hinge region

To explore a potential structural basis for the observed attenuation of NF-κB activity, 4-MC was docked using CB-dock3 (13) into the kinase domain of human IKKβ using chain B of PDB structure 4KIK, which represents the phosphorylated, catalytically active protomer of the asymmetric IKKβ dimer (12). The top-ranked pose produced an AutoDock Vina docking score of ™5.7 kcal/mol and was positioned within the canonical ATP-binding hinge region (Fig. 3A; Table 1).

**Table 1.** Predicted docking score and key residue interactions of 4-MC with IKKβ.

| Target | PDB ID | Docking score (kcal/mol) | Polar and hydrogen-bonding contacts | Hydrophobic contact(s) |
| --- | --- | --- | --- | --- |
| IKKβ (chain B) | 4KIK | -5.7 | Glu97, close O⋯O polar contact, 2.78 Å; Cys99, predicted bifurcated hydrogen-bonding contacts, 3.00 and 3.21 Å | Val29, Tyr98, Val152, Ile165 |

*Note: Interaction distances were measured from the top-ranked docked pose. The Glu97 interaction was classified as a close polar contact rather than a definitive hydrogen bond because the ligand hydroxyl-hydrogen orientation was not explicitly optimized or validated. Docking was performed against chain B of IKKβ (PDB: 4KIK) using CB-Dock3 with the AutoDock Vina scoring engine.*

**Fig. 3.**
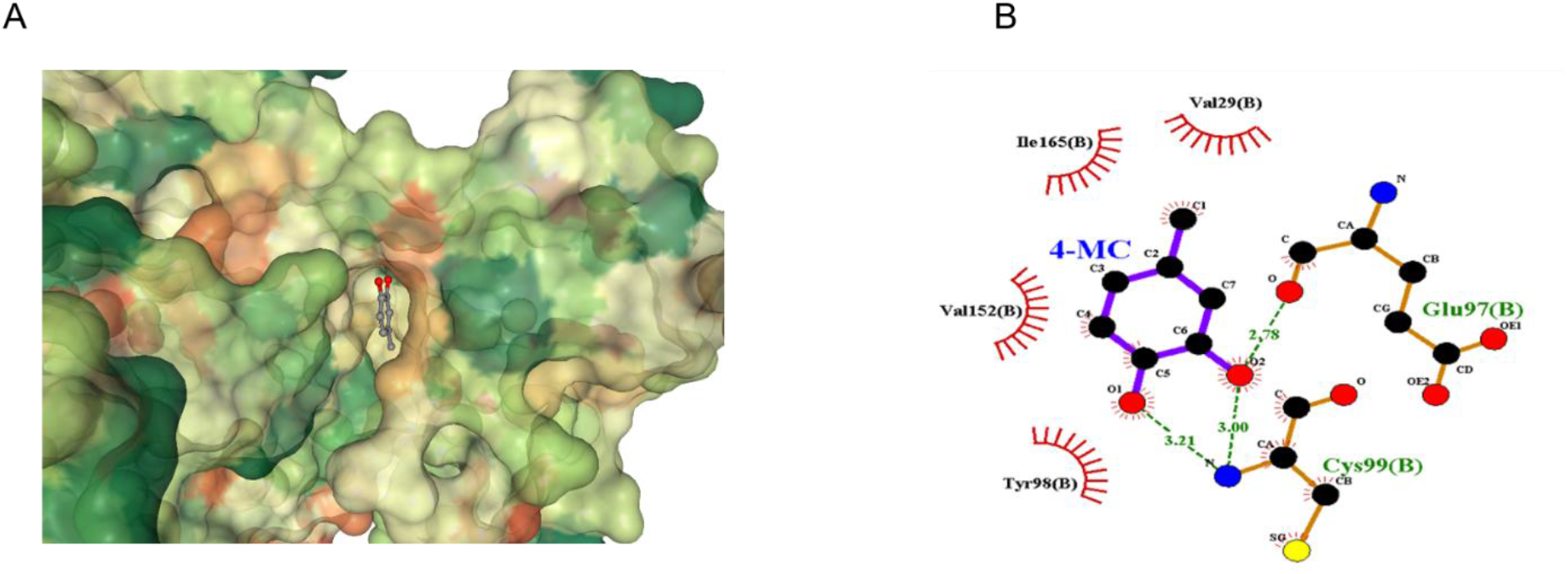
Predicted non-covalent docking pose of 4-methylcatechol within the IKKβ ATP-binding hinge region. Molecular docking was performed using chain B of human IKKβ from PDB structure 4KIK with CB-Dock3 and the AutoDock Vina scoring engine. **(A)** Surface representation showing 4-methylcatechol (4-MC) positioned within the ATP-binding cleft of IKKβ. **(B)** Two-dimensional interaction diagram generated using LigPlot+. The docked pose showed a close O···O polar contact with the backbone carbonyl oxygen of Glu97 at 2.78 Å and two predicted hydrogen-bonding contacts with the backbone amide nitrogen of Cys99 at 3.00 and 3.21 Å. Hydrophobic contacts were predicted with Val29, Tyr98, Val152, and Ile165.

In the predicted pose, the catechol hydroxyl groups of 4-MC were oriented toward the hinge backbone (Fig. 3B). One ligand oxygen was positioned 2.78 Å from the backbone carbonyl oxygen of Glu97. Because this represents an O···O contact and the orientation of the ligand hydroxyl hydrogen was not geometrically optimized, the interaction was conservatively classified as a close polar contact rather than a definitive hydrogen bond. The backbone amide nitrogen of Cys99 formed two predicted hydrogen-bonding contacts with the catechol oxygen atoms at distances of 3.00 and 3.21 Å, consistent with a bifurcated interaction. Hydrophobic contacts were additionally predicted with Val29, Tyr98, Val152, and Ile165.

Glu97, Tyr98, and Cys99 form part of the IKKβ hinge region engaged by ATP-site inhibitors, including the co-crystallized inhibitor K252a in the 4KIK structure. Thus, the predicted pose places 4-MC within a structurally established ligand-binding region of the IKKβ ATP pocket.

Owing to its small molecular size, 4-MC occupies only a limited portion of the ATP-binding cleft and may therefore exhibit fragment-like recognition of the hinge region. Together with the observed reduction in RANKL-induced NF-κB reporter activity, the docking results identify IKKβ as a plausible candidate target of 4-MC. However, the modest docking score and computational nature of the analysis should be considered. Docking alone cannot establish direct target engagement, kinase inhibition, or ATP-competitive activity, and these possibilities require biochemical or biophysical validation.

### 2.4. Covalent docking predicts engagement of oxidized 4-MC at Cys179 of IKKβ

Because 4-MC is an oxidizable catechol, it can generate the electrophilic quinone 4-methyl-1,2-benzoquinone (4MBQ) (14). Covalent docking was therefore performed to investigate whether 4MBQ could adopt a binding geometry compatible with covalent modification of IKKβ at Cys179, in addition to the non-covalent ATP-hinge interaction predicted for the parent 4-MC molecule. Cys179 is located within the IKKβ activation loop and has previously been implicated in the regulation of kinase activation (12) and in sensitivity to electrophilic inhibitors, including celastrol (15) and the synthetic triterpenoid CDDO-Me (16). CDDO and CDDO-Me form covalent adducts with wild-type IKKβ but not with an IKKβ C179A mutant, and CDDO-Me inhibits IKKβ activity by a mechanism dependent on oxidation of Cys179 (16).

Covalent docking was performed using GalaxyCDock (17) with ligand-embedded protein link-atom geometry incorporating the SG and CB atoms of Cys179. Five candidate models were generated and ranked using the GalaxyDock2-DL scoring function. The top-ranked model received a score of 21.851, while the five models had scores ranging from 21.851 to 23.236. These values were used only for ranking poses generated within the same docking run and do not represent binding free energies or values directly comparable with the AutoDock Vina docking score.

In the top-ranked model, 4MBQ formed a modeled covalent S–C linkage between the Cys179 side-chain sulfur and a quinone carbon atom (Fig. 4; Table 2). The modeled complex also showed a predicted polar contact with Thr200 and surrounding non-bonded contacts involving Ile142, Lys171, Leu173, Ser177, Leu178, Thr180, Tyr199, Val201, and Val203. The position of the predicted adduct near Ser177 and Ser181, the activation-loop phosphorylation sites associated with IKKβ activation, provides a structurally plausible context in which modification of Cys179 could influence activation-loop organization. Nevertheless, proximity to these regulatory residues does not by itself demonstrate an effect on phosphorylation or catalytic activity.

**Table 2.** Summary of covalent docking of oxidized 4-MC (4MBQ) at Cys179 of IKKβ.

| Target | Reactive residue | Docking method | Best model score (GalaxyDock2-DL) | Key interactions |
| --- | --- | --- | --- | --- |
| IKKβ (chain B) | Cys179 | GalaxyCDock with ligand-embedded Cys179 SG/CB link-atom geometry | 21.851 (Model 1 of 5) | Modeled Cys179-SG–C(4MBQ) covalent linkage; predicted polar contact with Thr200; non-bonded contacts with Ile142, Lys171, Leu173, Ser177, Leu178, Thr180, Tyr199, Val201, and Val203 |

*Note: Docking was performed using GalaxyCDock with the ligand-embedded protein link-atom geometry option, incorporating the SG and CB atoms of Cys179. Scores are reported as GalaxyDock2-DL values, for which lower values indicate more favorable predicted binding geometry. The newly modeled protein–ligand bond is between Cys179 SG and a carbon atom of 4MBQ; the SG–CB bond represents the native Cys179 side-chain connectivity.*

**Fig. 4.**
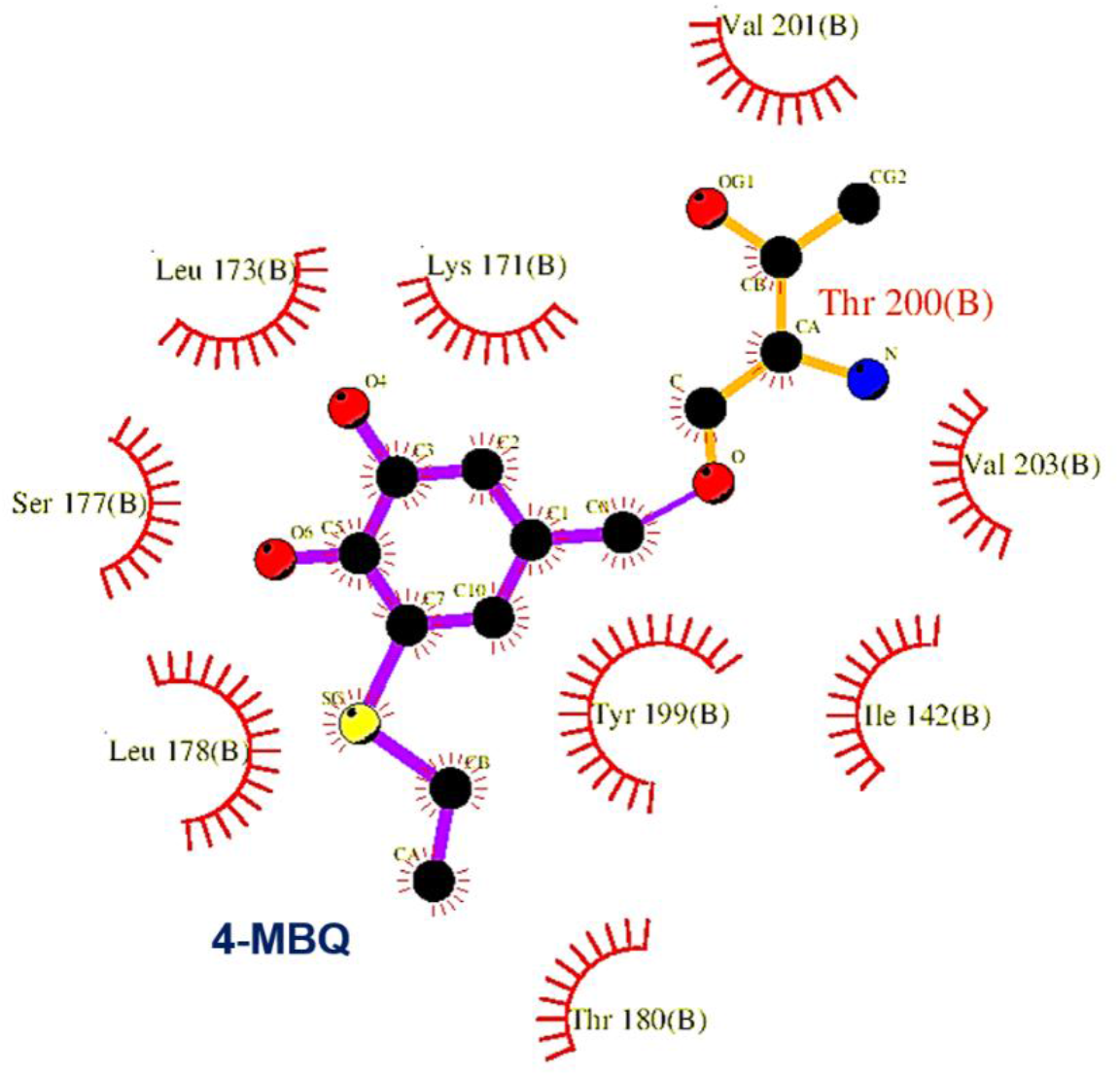
Predicted covalent binding mode of oxidized 4-MC (4MBQ) at Cys179 of IKKβ. Covalent docking of oxidized 4-methylcatechol, 4-methyl-1,2-benzoquinone (4MBQ), was performed against Cys179 of IKKβ chain B using GalaxyCDock with ligand-embedded protein link-atom geometry. The two-dimensional interaction diagram of the top-ranked model was generated using LigPlot+. The model contains a covalent S–C linkage between the Cys179 side-chain sulfur and a carbon atom of 4MBQ, a predicted polar contact with Thr200, and surrounding non-bonded contacts involving Ile142, Lys171, Leu173, Ser177, Leu178, Thr180, Tyr199, Val201, and Val203. The SG and CB atoms of Cys179 were incorporated into the ligand-embedded docking input and are therefore depicted as part of the ligand structure rather than as a separately labeled residue in the LigPlot+diagram.

Recent mechanistic analysis of solution-phase thiol addition to 4-methyl-ortho-benzoquinone indicates that adduct formation may proceed through a free-radical chain pathway rather than exclusively through a conventional polar 1,4-addition mechanism (14). Covalent docking does not simulate electron transfer, thiyl-radical generation, reaction kinetics, or the chemical mechanism of bond formation. The present analysis therefore supports only the geometric plausibility of the post-reaction 4MBQ–Cys179 adduct and does not establish how or whether the reaction occurs under cellular conditions.

The predicted cysteine-reactive behavior of 4MBQ may also be considered in relation to the previously reported effects of 4-MC on Keap1/Nrf2 signaling (11). Keap1 functions as an electrophile- and redox-sensitive regulator whose activity can be altered through modification of reactive cysteine residues. However, the reported modulation of Keap1/Nrf2 signaling by 4-MC does not demonstrate that 4MBQ directly forms covalent adducts with Keap1. It is therefore plausible, but currently unproven, that oxidation of 4-MC could enable interactions with multiple cysteine-containing regulatory proteins. Keap1/Nrf2/HO-1 activation and Cys179-dependent IKKβ modification should consequently be regarded as potentially concurrent but experimentally unresolved mechanisms.

Overall, the covalent docking model identifies Cys179 as a chemically and structurally plausible candidate site for quinone-mediated engagement of IKKβ. However, the analysis does not demonstrate intracellular formation of 4MBQ, direct target engagement, site-selective Cys179 modification, or inhibition of IKKβ kinase activity. Confirmation would require experiments such as intact-protein or peptide-level mass spectrometry, comparison with an IKKβ C179A mutant, direct kinase assays, or cellular target-engagement analysis.

### 2.5. Quantum-chemical analysis supports enhanced electrophilicity of oxidized 4-MC

To examine how oxidation alters the electronic reactivity of 4-MC, frontier molecular orbital properties of the parent catechol and its oxidized quinone, 4-methyl-1,2-benzoquinone (4MBQ), were calculated at the B3LYP (18, 19)/6-31G(d) (20) level using PySCF (21). Compared with 4-MC, 4MBQ exhibited a substantially lower LUMO energy and a narrower HOMO–LUMO gap (3.272 versus 5.912 eV; Table 3). The global electrophilicity index also increased approximately sixfold following oxidation, from 1.316 eV for 4-MC to 7.869 eV for 4MBQ.

**Table 3.**
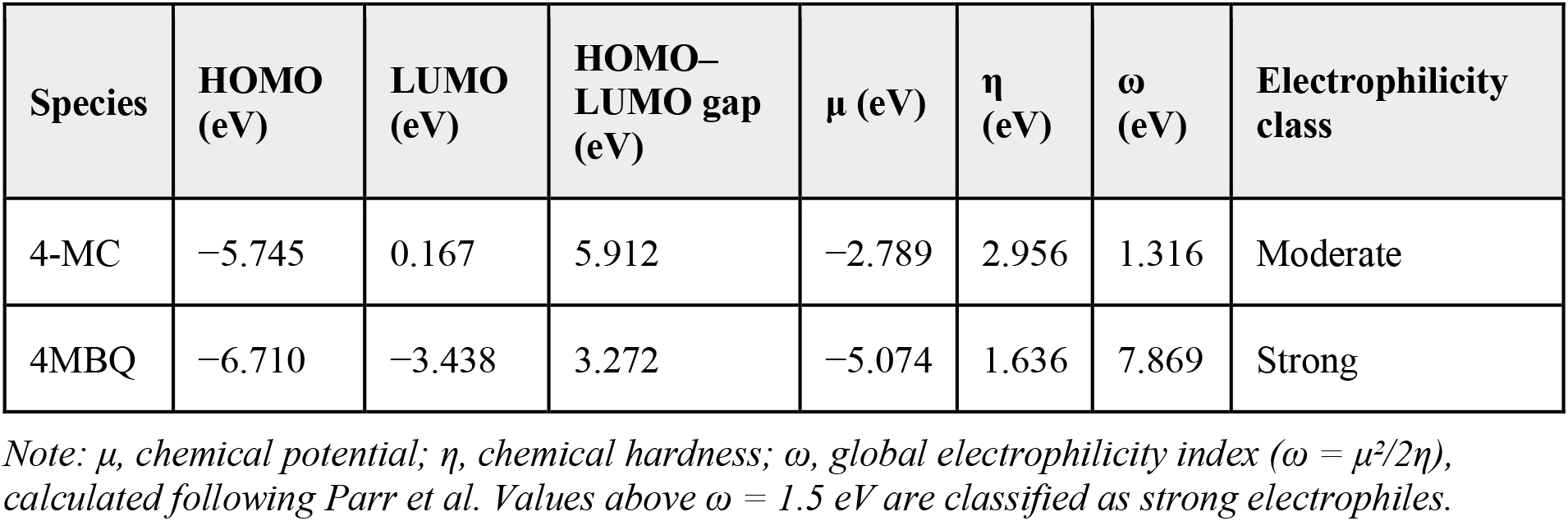
Frontier molecular orbital energies and electrophilicity indices of 4-MC and its oxidized quinone, 4MBQ.

| Species | HOMO (eV) | LUMO (eV) | HOMO–LUMO gap (eV) | $\mu$ (eV) | $\eta$ (eV) | $\omega$ (eV) | Electrophilicity class |
| --- | --- | --- | --- | --- | --- | --- | --- |
| 4-MC | –5.745 | 0.167 | 5.912 | –2.789 | 2.956 | 1.316 | Moderate |
| 4MBQ | –6.710 | –3.438 | 3.272 | –5.074 | 1.636 | 7.869 | Strong |

*Note: $\mu$ , chemical potential; $\eta$ , chemical hardness; $\omega$ , global electrophilicity index ( $\omega = \mu^2/2\eta$ ), calculated following Parr et al. Values above $\omega = 1.5$ eV are classified as strong electrophiles.*

The global electrophilicity index, ω, was calculated from the electronic chemical potential and chemical hardness using the conceptual DFT definition introduced by Parr and co-workers (22). According to the empirical electrophilicity scale subsequently proposed by Domingo and co-workers, compounds with ω > 1.5 eV are classified as strong electrophiles, whereas those with values of 0.8– 1.5 eV are classified as moderate electrophiles (23). Accordingly, 4MBQ was classified as a strong electrophile, while the parent 4-MC molecule fell within the moderate-electrophile range.

Frontier-orbital visualization showed that the low-lying LUMO of 4MBQ was distributed across the conjugated quinone π-system, including the carbonyl and ring regions (Fig. 5). This distribution, together with the markedly lower LUMO energy and increased electrophilicity index, is consistent with enhanced electron-accepting character following oxidation. However, the global electrophilicity index and orbital isosurfaces do not independently identify the preferred site or chemical mechanism of thiol addition. In particular, mechanistic studies of thiol addition to 4-methyl-*ortho*-benzoquinone support a thiyl-radical chain pathway rather than a simple classical nucleophilic 1,4-addition mechanism (14).

**Fig. 5.**
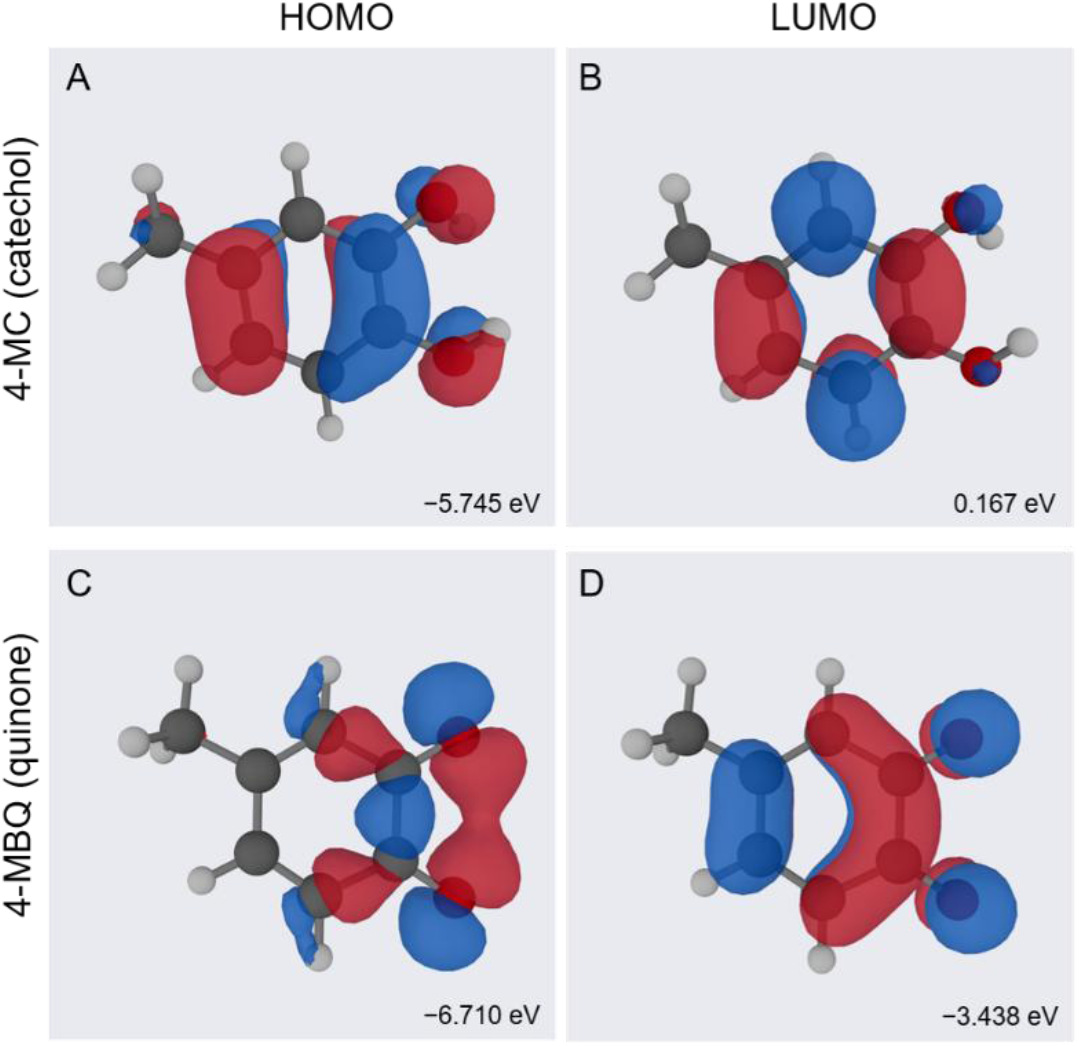
Frontier molecular orbitals of 4-methylcatechol and its oxidized quinone. HOMO and LUMO isosurfaces of 4-methylcatechol (4-MC) and 4-methyl-1,2-benzoquinone (4MBQ) were calculated at the B3LYP/6-31G(d) level and visualized using MOrbVis. **(A)** HOMO of 4-MC. **(B)** LUMO of 4-MC. **(C)** HOMO of 4MBQ. **(D)** LUMO of 4MBQ. Orbital energies are indicated in electronvolts below the respective structures. Red and blue surfaces represent opposite phases of the molecular orbitals and do not represent positive and negative atomic charges.

Overall, these calculations indicate that oxidation of 4-MC to 4MBQ markedly enhances its electrophilic character and produces an electronic structure compatible with covalent thiol engagement. The results therefore complement the predicted 4MBQ–Cys179 covalent docking model. Nevertheless, they do not demonstrate intracellular formation of 4MBQ, reaction with Cys179, site selectivity, or the mechanism and kinetics of covalent-bond formation.

### 2.6. In silico pharmacokinetic, safety, and medicinal-chemistry profile of 4-MC

The predicted physicochemical, pharmacokinetic, toxicity, and medicinal-chemistry properties of 4-MC were evaluated using SwissADME (24) and pkCSM (25) (Table 4). 4-MC satisfied Lipinski’s rule of five without any violations. SwissADME classified the compound as having high gastrointestinal absorption, while pkCSM predicted an intestinal absorption of 91.7%. The compound was also predicted to cross the blood–brain barrier. Although these predictions suggest favorable passive absorption and distribution characteristics, blood–brain barrier permeability is not necessarily advantageous for a compound intended to act primarily on peripheral bone-resorptive pathways.

**Table 4.** Predicted ADMET and drug-likeness properties of 4-MC.

| Property | Predicted value | Source |
| --- | --- | --- |
| Molecular weight | 124.14 g/mol | SwissADME |
| Lipinski violations | 0 | SwissADME |
| GI absorption | High (91.7%) | pkCSM |
| BBB permeant | Yes | SwissADME / pkCSM |
| CYP450 substrate/inhibitor (5 isoforms) | None predicted | pkCSM |
| AMES mutagenicity | Negative | pkCSM |
| Hepatotoxicity | Negative | pkCSM |
| hERG I/II inhibition | Negative | pkCSM |
| Skin sensitization | Positive | pkCSM |
| PAINS / Brenk structural alerts | Catechol alert (both filters) | SwissADME |

No substrate or inhibitory liabilities were predicted among the evaluated major cytochrome P450 endpoints. In addition, 4-MC was predicted to be negative for Ames mutagenicity, hepatotoxicity, and inhibition of hERG I and II channels. These results indicate a generally favorable preliminary computational profile; however, they should not be interpreted as evidence of experimental safety or metabolic stability.

Potential liabilities were also identified. 4-MC was predicted to cause skin sensitization and generated PAINS and Brenk alerts because of its catechol substructure. These filters identify structural features associated with possible assay interference, redox activity, chemical instability, or nonspecific reactivity, but they do not establish that such effects occur under the experimental conditions used here. Nevertheless, the alerts are chemically compatible with the ability of catechols to undergo oxidation to electrophilic quinones and therefore warrant consideration when interpreting the biological and docking results.

Future studies should incorporate orthogonal controls to distinguish pathway-specific activity from redox-dependent or chemically reactive effects. These could include direct analysis of 4-MC oxidation, glutathione- or cysteine-adduct formation by mass spectrometry, comparison under controlled reducing and nonreducing conditions, and evaluation of whether thiol-containing scavengers alter its activity. Such experiments would be more informative than relying solely on PAINS or toxicity predictions to infer the mechanism of action.

### 2.7. Limitations and future directions

Several limitations should be acknowledged. NF-κB reporter activity was evaluated at only one concentration of 4-MC, and concentration–response analysis is therefore needed to define its potency in this system. Additional osteoclast markers, including NFATc1, c-Fos, cathepsin K, and DC-STAMP, together with bone-resorption assays, would further characterize its anti-osteoclastogenic effects.

The predicted non-covalent interaction of 4-MC within the IKKβ ATP-binding site and covalent engagement of Cys179 by 4MBQ remain computational hypotheses. Direct kinase and target-engagement assays are required to evaluate the non-covalent mechanism, whereas mass spectrometry and comparison of wild-type IKKβ with a C179A or C179S mutant would help confirm covalent modification. The intracellular formation of 4MBQ and its thiol adducts should also be examined under defined redox conditions. Finally, validation in primary osteoclast precursors and appropriate animal models will be required to establish the physiological and translational relevance of these findings.

### 2.8. Conclusion

In conclusion, 4-MC suppresses RANKL-induced NF-κB reporter activity and inhibits RANKL-induced TRAP activity and osteoclast differentiation in a concentration-dependent manner (IC_50_ = 2.15 µM), without reducing cell viability across the concentrations tested. Molecular docking suggests a possible interaction with the ATP-binding hinge region of IKKβ, and computational analyses further raise the possibility of covalent engagement of Cys179 by an oxidized quinone metabolite of 4-MC. These findings identify 4-MC as a small catechol scaffold worthy of further mechanistic and structure-activity investigation.

## 3. Experimental

### 3.1. Reagents and compounds

4-Methylcatechol (4-MC) was obtained from MedChemExpress (CAS No.: 452-86-8; Catalog no. HY-W012814; purity ≥99%). A 10 mM stock solution prepared in dimethyl sulfoxide (DMSO) was stored at ™80°C. Working solutions were freshly prepared by dilution in culture medium. The final DMSO concentration did not exceed 0.1% (v/v) in any experiment. Recombinant murine RANKL was purchased from R&D Systems (catalog no. 462-TEC-010). Cell Counting Kit-8 (CCK-8) reagent was obtained from ServiceBio.

### 3.2. Cell culture

RAW264.7 murine macrophage cells were cultured in Dulbecco’s modified Eagle’s medium supplemented with 10% fetal bovine serum and 1% penicillin-streptomycin. Cells were maintained at 37°C in a humidified atmosphere containing 5% CO2 as described earlier (26, 27).

### 3.3. TRAP activity assay

RAW264.7 cells were seeded in 96-well plates at 3 × 10^3^ cells per well. After overnight attachment, the cells were treated with RANKL at 50 ng/mL in the presence of a dilution series of 4-MC ranging from 1 to 10 µM, or in the presence or absence of 4-MC at a fixed concentration of 2 µM for osteoclast staining and quantification. Treatment was continued for 3 days, and the culture medium was replaced every 48 hours. Tartrate-resistant acid phosphatase (TRAP) activity was measured using a p-nitrophenyl phosphate-based colorimetric assay. Cells were lysed with 0.1% Triton X-100. The lysates were incubated at 37°C with acetate buffer containing p-nitrophenyl phosphate and sodium tartrate. The release of p-nitrophenol was measured at 405 nm using a microplate reader. TRAP activity was normalized to the vehicle control, and percent inhibition at each 4-MC concentration was calculated relative to the RANKL-alone control. The half-maximal inhibitory concentration (IC_50_) was determined by nonlinear regression (log[inhibitor] vs. normalized response) using GraphPad Prism version 8.0. For osteoclast staining and quantification, RAW264.7 cells were treated with RANKL (50 ng/mL) in the presence or absence of 4-MC (2 µM) for 3 days, fixed, and stained using a TRAP staining kit according to the manufacturer’s instructions (ServiceBio). TRAP-positive multinucleated cells containing three or more nuclei were counted as osteoclasts under a light microscope, and the number of multinucleated osteoclasts per well was quantified across multiple fields.

### 3.4. NF-κB luciferase reporter assay

HEK-293T cells stably expressing an NF-κB-responsive firefly luciferase reporter (NFκB-TA-Luc-EF1α-mCherry, Beyotime) and RANK (Mouse TNFRSF11A, Sino Biological) were used. Cells were seeded in 96-well plates at a density of 1 × 10^4^ cells per well. After overnight attachment, cells were pretreated with 1 µM 4-MC for 1 h, followed by stimulation with 50 ng/mL RANKL for 6 h. Luciferase activity was measured using the Steady-Glo Luciferase Assay System (Promega) according to the manufacturer’s instructions. Luminescence values were normalized to vehicle control and expressed as relative fold change.

### 3.5. CCK-8 cell viability assay

RAW264.7 cells were seeded in 96-well plates at 5 × 10^3^ cells per well. After overnight attachment, the cells were treated with 4-MC, or an equivalent volume of vehicle, for 48 hours. CCK-8 assay was then performed as per manufacturer’s protocol (ServiceBio, China). Cell viability was expressed as a percentage of the vehicle control. Results are presented in the Supporting information (Fig. S1, S2).

### 3.6. Molecular docking

Molecular docking was performed using the CB-Dock3 web server (13). The platform combines automated cavity detection with AutoDock Vina-based molecular docking and scoring. The crystal structure of human IKKβ was obtained from the Protein Data Bank under accession number 4KIK (12). The structure contains an asymmetric IKKβ dimer with two protomers in different conformational states. Chain B was selected for docking because it represents the phosphorylated protomer in the active kinase conformation. The co-crystallized inhibitor K252a and crystallographic water molecules were removed before docking. The three-dimensional structure of 4-MC was prepared and submitted to CB-Dock3 for blind cavity detection and docking against chain B. The highest-ranked pose located within the canonical ATP-binding pocket was selected for interaction analysis. Two-dimensional protein-ligand interaction diagrams were generated using LigPlot+ (28). Hydrogen-bonding interactions were identified using a donor-acceptor distance cutoff of less than 3.5 Å.

### 3.7. Covalent docking analysis

Because 4-MC contains a catechol moiety that can undergo oxidation to an electrophilic ortho-quinone (4-methyl-ortho-benzoquinone, 4MBQ), covalent docking was performed to evaluate whether the oxidized form of 4-MC could engage IKKβ through covalent modification of a reactive cysteine. Covalent docking was performed using the GalaxyCDock web server (17). The reactive ring carbon of 4MBQ was assigned based on the reported regiochemistry of thiol addition to 4-methyl-ortho-benzoquinone, in which the major addition product (the 5-S-adduct) forms at the ring position distal to the alkyl substituent (14). Cys179, a residue within the IKKβ activation loop previously shown to be covalently targeted by structurally unrelated electrophilic natural products including celastrol and the methyl ester of CDDO, was selected as the candidate reactive residue. Docking was performed using the ligand-embedded protein link-atom option, with the ligand 4MBQ modified to include SG and CB link atoms corresponding to the Cys179 side chain. The top-ranked covalent binding mode was visualized using LigPlot+ (28).

### 3.8. Quantum chemical reactivity analysis

To evaluate the intrinsic electrophilic reactivity conferred by oxidation of 4-MC to its ortho-quinone form, ground-state geometries of 4-MC and 4MBQ were generated using the MMFF94 force field in RDKit and subjected to single-point density functional theory (DFT) calculations at the B3LYP/6-31G(d) level using PySCF. Frontier molecular orbital energies (highest occupied molecular orbital, HOMO; lowest unoccupied molecular orbital, LUMO) were used to calculate the chemical potential (μ = (E_HOMO + E_LUMO)/2), chemical hardness (η = (E_LUMO ™ E_HOMO)/2), and global electrophilicity index (ω = μ^2^/2η), following the formalism of Parr et al. (22).

### 3.9. In silico ADMET and drug-likeness prediction

Physicochemical properties and drug-likeness of 4-MC were evaluated using SwissADME (24). Absorption, distribution, metabolism, excretion, and toxicity (ADMET) properties, including gastrointestinal absorption, blood-brain barrier permeability, cytochrome P450 substrate and inhibitor status, and predicted toxicity endpoints (including Ames mutagenicity, hepatotoxicity, hERG inhibition, and skin sensitization), were evaluated using pkCSM (25).

### 3.10. Statistical analysis

Data are presented as the mean ± standard deviation from at least three independent experiments. Comparisons among the three or more groups were performed using one-way analysis of variance followed by Tukey’s multiple-comparisons test. Comparisons between two groups were performed using an unpaired two-tailed Student’s t-test. Half-maximal inhibitory concentration (IC50) values were calculated by nonlinear regression (log[inhibitor] vs. normalized response). Statistical analysis was performed using GraphPad Prism version 8.0 (GraphPad Software, San Diego, CA, USA). A value of p < 0.05 was considered statistically significant.

## CRediT authorship contribution statement

**Vishwa Deepak:** Conceptualization, Supervision, Methodology, Funding acquisition, Investigation, Writing - review & editing. **Chengxu Xie:** Investigation, Writing - original draft. **Lifang Zhang:** Investigation, Writing - original draft. **Xinyi Bao:** Investigation, Writing - original draft. **Xiaohan Li:** Investigation, Writing - original draft. **Yili Ding:** Resources, Methodology, Writing - review & editing. **Mojtaba Tabandeh:** Resources, Methodology, Writing - review & editing. **Farwa Basit:** Resources, Methodology, Writing - review & editing. **Heriberto Velez:** Resources, Methodology, Writing - review & editing. **Santosh Kumar:** Resources, Methodology, Writing - review & editing.

## Declaration of Competing Interest

The authors declare that they have no known competing financial interests or personal relationships that could have appeared to influence the work reported in this paper.

## Funding

This research was funded by the WKU-ISRG grant (ISRG2023032) and the International Frontier Interdisciplinary Research Institute (IFIRI) of Wenzhou-Kean University grant (KY20250604000450) awarded to Vishwa Deepak. Additional support was provided by the Wenzhou-Kean University Talent Introduction Grant (WB20220901000092), awarded to Yili Ding and by the WKU-ISRG grant (ISRG2023027) awarded to Mojtaba Tabandeh.

## Data availability

The authors confirm that all data generated during this study are included in this published article and its supplementary information.

## Institutional review board statement

Not applicable.

## Informed consent statement

Not applicable.

## Supporting information

Figure S1: 4-MC does not affect HEK-293T/RANK NF-κB luciferase reporter cell viability.

Figure S2: 4-MC does not affect RAW264.7 cell viability.

## Notes

### Competing Interest Statement

The authors have declared no competing interest.

