## Supplementary material for "IKKβ as a putative non-covalent and quinone-mediated covalent target of 4-methylcatechol in RANKL/NF-κB signaling: a combined computational and experimental analysis": Fig. S1, S2

### Supplementary Information

**Figure S1. 4-MC does not affect viability of HEK-293T/RANK NF-κB luciferase reporter cells.**

HEK-293T/RANK NF-κB luciferase reporter cells were treated with 4-MC at 1 μM, or with vehicle, for 48 hours. Cell viability was measured using the CCK-8 assay and expressed as a percentage of the vehicle control. Data are presented as the mean ± SD of at least three independent experiments. Statistical analysis was performed using an unpaired two-tailed Student's t-test. ns, not significant.

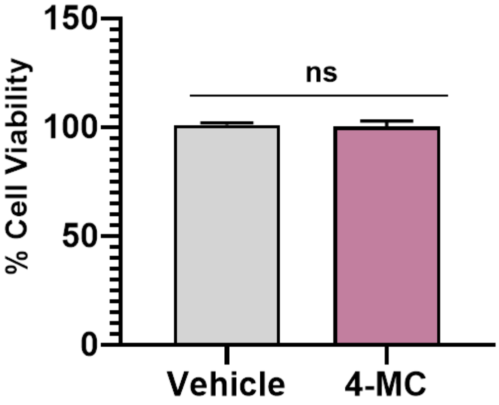

**Figure S2. 4-MC does not affect RAW264.7 cell viability.**

RAW264.7 cells were treated with 4-MC at the indicated concentrations or with vehicle for 48 hours. Cell viability was measured using the CCK-8 assay and expressed as a percentage of the vehicle control. Data are presented as the mean ± SD of at least three independent experiments. Statistical analysis was performed using one-way ANOVA followed by Tukey's multiple-comparisons test. ns, not significant.

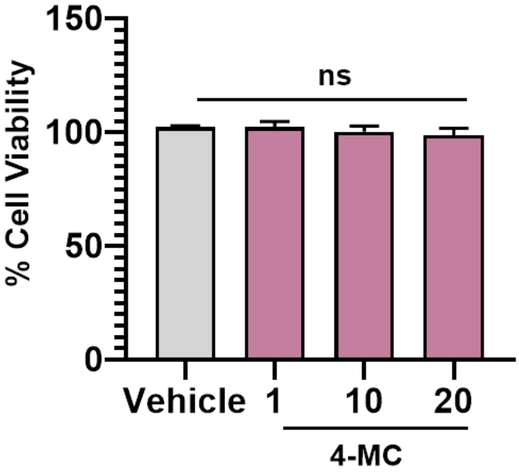
